# Conserved conformational hierarchy across functionally divergent glycosyltransferases of the GT-B structural superfamily as determined from microsecond molecular dynamics

**DOI:** 10.1101/2020.07.02.181073

**Authors:** Carlos A. Ramirez-Mondragon, Megin E. Nguyen, Jozafina Milicaj, Frank J. Tucci, Ramaiah Muthyala, Erika A. Taylor, Yuk Y. Sham

**Affiliations:** Department of Chemistry, Wesleyan University, Middletown, CT 06459, United States; Department of Integrative Biology and Physiology, Medical School, University of Minnesota, Minneapolis, MN 55455, United States; Department of Experimental and Clinical Pharmacology, College Pharmacy, University of Minnesota, Minneapolis, MN 55455, United States; Bioinformatics and Computational Biology Program, University of Minnesota

## Abstract

It has long been understood that some proteins to undergo conformational transitions enroute to the Michaelis Complex to allow chemistry. Examination of crystal structures of glycosyltransferase enzymes in the GT-B structural class reveals that the presence of ligand in the active site is necessary for the protein to crystalize in the closed conformation. Herein we describe microsecond molecular dynamics simulations of two evolutionarily unrelated glycosytransferases, HepI and GtfA. Simulations were performed using these proteins in the open and closed conformations, (respectively,) and we sought to identify the major dynamical modes and communication networks which allow conformational transition between the open and closed structures. We provide the first reported evidence (within the scope of our experimental parameters) that conformational hierarchy/directionality of the interconversion between open and closed conformations is a conserved feature of enzymes of the same structural superfamily. Additionally, residues previously identified to be important for substrate binding in HepI were shown to have strong negative correlations with non-ionizable residues distal to the active site. Mutagenesis of these residues produced mutants with altered enzymatic efficiency exhibiting lower K_m_ values, while the k_cat_ is effectively unchanged. The negatively correlated motions of these residues are important for substrate binding and forming the Michaelis complex, without impacting the activation barrier for catalysis. This suggests that in the bi-domain HepI, the enzyme dynamics did not impact the transition state stabilization or chemistry, but rather earlier steps along the reaction coordinate, leading to the reorganization of the active site electrostatic environment required for catalysis.

## Introduction

Glycosylation is a highly regulated, ubiquitous biochemical process catalyzed by glycosyltransferase (GT, E.C. 2.4.x.y) enzymes. The implications of glycosylation in cellular processes are broad, as it modulates the structure, stability, and hence function of the target molecule. Although GTs constitute approximately 1 to 2% of the genomes that have been sequenced from species across living kingdoms^1^, the details of the molecular mechanism of many GTs remain relatively undetermined. Significant research endeavors have identified and characterized key residues and regions linked to the enzymatic cycle in diverse GTs.^2-9^ The data gathered from these investigations is furthering the development of an atomistic description of the molecular mechanism in GTs, which is paramount for inhibitor discovery, chimeric protein design, enzymatic control and regulation, among other studies that offer potentially beneficial clinical and industrial applications.

GTs mediate the transfer of a single sugar (monosaccharide) from an activated sugar complex (co-substrate) onto a select acceptor substrate.^10^ The identities of the substrates and co-substrates are GT specific, with the pool of available co-substrates encompassing a myriad of monosaccharide and activation moiety conjugates (with the latter including nucleotide-monophosphate, nucleotide-diphosphate, lipid-phosphate, or unsubstituted-phosphate compounds).^10^ The ODLA acceptor substrates range from single biomolecules to macromolecular complexes of varying composition (e.g., monosaccharides, oligosaccharides, organic molecules, lipids, polypeptides, DNA).^10^ The resulting sugar conjugate product, in most reported cases, serves as the preferred substrate for a subsequent GT in a series of glycosylation reactions that lead to the final multi-glycosylated product.^3-5, 11^

The largest subset of identified GTs in CAZy (Carbohydrate-active enzymes database) correspond to the bi-domain Leloir-GTs (monosaccharide-nucleotide dependent).^12^ The individual domains of Leloir-GTs are named according to their relative primary sequence composition (*N* and *C* domain) while their combined spatial configuration determines the structural superfamily (GT-A or GT-B).^13^ Regardless of the structural superfamily classification, Leloir-GT structures depict the nucleotide binding Rossman fold motif (βαβ) as a conserved feature that comprises the core of all resolved domains. GT-Bs such as Heptosyltransferase I (HepI) and Glycosyltransferase A (GtfA) consist of topologically identical domains joined by an extended linker (loop-α-loop) region that are positioned in a stacked configuration (**Fig. 1**). This arrangement generates a central inter-domain cavity that serves as the binding site for both ligands and contains the enzyme’s catalytic core. The binding position of each ligand in the GT-B cavity is domain specific, as the *N* and *C* domain demonstrate selective binding affinity towards the substrate and co-substrate, respectively.^3-5^ The orientation of the ligands in the binding pocket further divides the cavity into catalytic (proximal to active site residues) and non-catalytic regions. GTs are further classified mechanistically as either retaining or inverting.^10^ The classification is given by comparing the stereochemistry of the newly formed glycosidic linkage of the product to that of the starting co-substrate.

**Figure 1.**
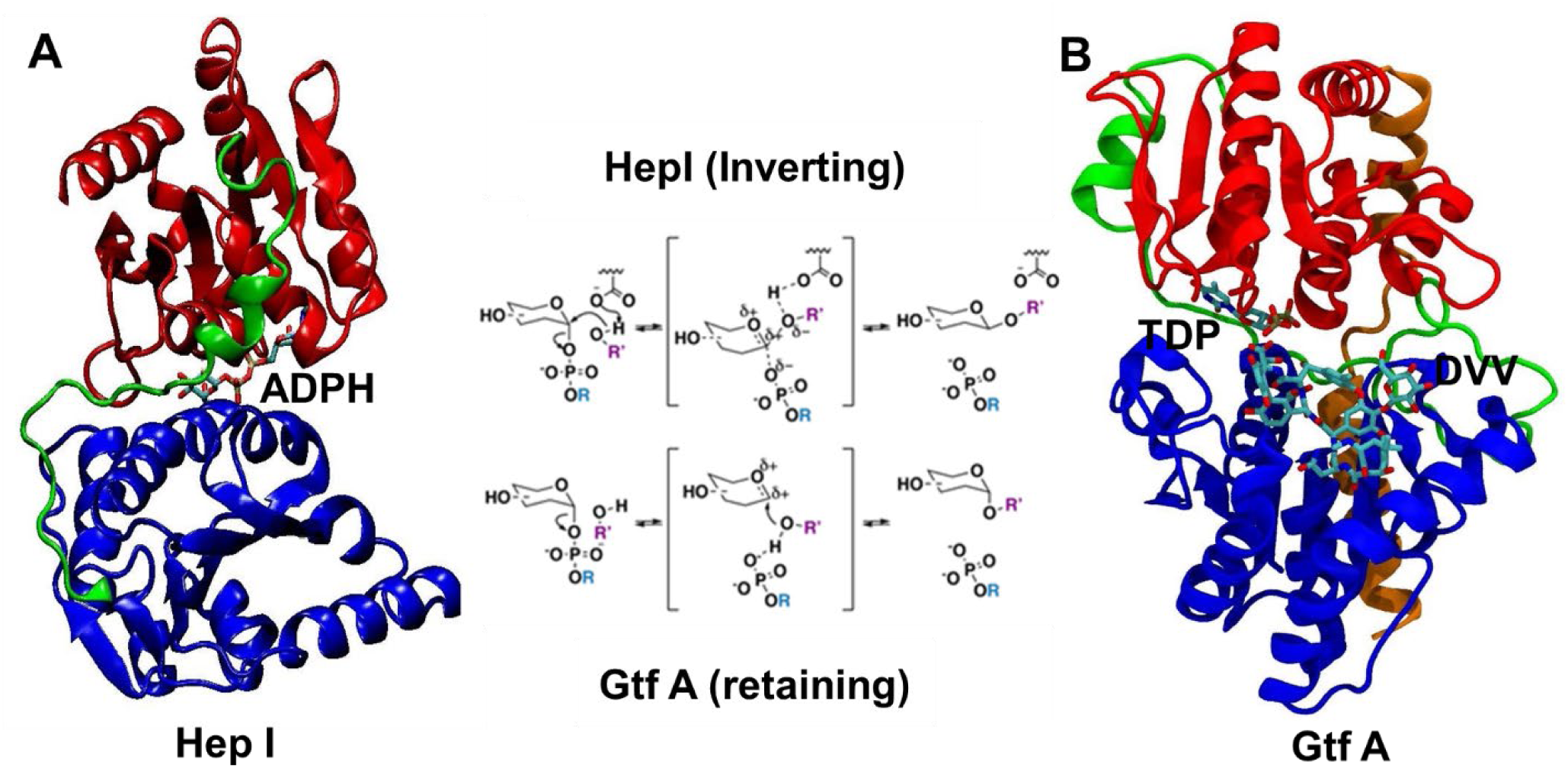
Ribbon representation of **A)** HepI (PDB: 2H1H) and **B)** GtfA (PDB: 1PN3) colored by N domain, C domain, linker, and spine (only in GtfA) region (blue, red, green, orange, respectively). HepI and GtfA co-factors, ADPH and TDP, respectively, and GtfA substrate DVV, are shown as ball and stick representation between N and C domains. The Center panel shows the inverting and retaining mechanism of glycosyltransfer reaction catalyzed by HepI and GtfA, respectively.

As observed experimentally, the transition from a catalytically inactive (open, apo) to active (closed, ternary) state in GT-Bs is induced by either substrate or co-substrate binding and is proposed to proceed via domain motions whose axis of rotation/translation is centered on residues of the linker and/or the extended C-terminal spine region (if present).^4-5, 11, 14^ These motions are accompanied by different degrees of peripheral α-helix repositioning and loop restructuring with no significant alteration to the underlying Rossman fold structure of each domain.^4-5^ The end result of this dynamic process, described here as domain flexibility, positions the domains closer to one another, emphasizing pre-existing or creating new intra- and inter-domain contacts while bridging the distance between the enzymatic and ligand reactive centers.^4-5^ Crystallographic examples of domain flexibility in GT-Bs include Gram-positive bacterial enzymes Mycothiol A (MshA, retaining, EC: 2.4.1.250), which mediates the first step of mycothiol biosynthesis^4^, and Glycosyltransferase A (GtfA, inverting, EC: 2.4.1.311), involved in biosynthesis of the natural glycopeptide antibiotic chloroeremomycin.^5^ Further evidence of conformational changes in GT-Bs upon substrate binding comes from intrinsic tryptophan fluorescence (ITF) experiments of the Gram-negative bacterial enzyme Heptosyltransferase I (HepI, inverting, EC: 2.4.99.B6).^15^ HepI catalyzes the addition of the first monosaccharide to the nascent polysaccharide core of the outer membrane lipopolysaccharide (LPS).^16-19^ Sole addition of a Heptose sugar to the acceptor substrate (O-deacylated *E. coli* Kdo2-lipid A, ODLA) analog resulted in a spectral blue-shift of the enzyme’s fluorescence profile. Such a shift is indicative of associated structural changes that desolvate one or more Trp residues. The extent and degree of the HepI motion cannot be described in the ensemble averaged fluorescence experiments.

Given the evolutionary conserved structure, chemical mechanism, and pre-requisite domain flexibility transitions required for function across inverting GT-Bs, we began to explore whether conformational dynamics is also a conserved feature of these familial enzymes. To do so, we characterize the type, degree, and order of the holo to apo transition observed domain flexibility motions in HepI and GtfA as reported by microsecond molecular dynamics (MD) trajectories generated on the Anton supercomputer.^20^ The advent of Anton has allowed for the atomistic examination of the dynamic behavior and associated properties of large biomolecular systems in explicit solvent at timescales that range from microseconds to early milliseconds.^21-24^ The selection of the GT-B’s GtfA and HepI for MD study pertains to: 1) the availability of high resolution holo-form structures with no unresolved back-bone or C_α_ atom coordinates; 2) a shared inversion catalytic mechanism; and 3) a common catalytic core, discovered during this study, which has been evolutionary conserved in both sequence and three-dimensional (3-D) space despite the divergent function of each enzyme.

## Methods

### Multiple Sequence Alignment (MSA)

The MSA was generated by filtering the HepI (UniProtKB: P24173) and GtfA (UniProtKB: P96558) sequences on the European Bioinformatics Institutes (EMBLEBI) Clustal Omega MSA web server with default settings. The resulting MSA was visualized and sequence identity conservation was determined using the Unipro UGENE (v. 1.14.2) bioinformatics software.

### Molecular Dynamics (MD) Simulations

Protein systems are built and equilibrated from their respective crystal structure coordinates (PDB Databank) using the CHARMM MD simulation program (v. c38a2) and the CHARMM c36 all-atom force field with CMAP corrections. All selected crystal structures met the following criteria: resolution ≤ 2.5 Å, R-Factor ≤ 0.2, and R-Free ≤ 0.30. Missing atoms and hydrogens were built using CHARMM. Proper protonation states of charged residues were investigated by means of calculating the probable pK_a_ (H++ server) for said residue and by exhaustive visual examination of the surrounding environment. For histidine residues, protonation was assigned on the basis of what proton position (δ or ε) generated the most favorable hydrogen bonding network. Apo systems were set up by ligand exclusion from the respective open-binary (HepI, PDB: 2H1H: Chain A) and closed-ternary (GtfA, PDB:1PN3: Chain A) structures. Both systems were solvated using the TIP3P explicit water model in a cubic box via the MMTSB toolset. Crystal water positions were retained, and depending on the system, sodium or chloride counter-ions were added to ensure electrostatic neutrality. SHAKE algorithm was applied to all bonds involving hydrogen. Improper contacts were handled via minimization with fixed restraints on all protein backbone, ligand, and crystal water heavy atoms using 25-50 steps of Steepest Descent minimization followed by 25-50 steps of adopted basis Newton-Raphson Method minimization. Careful consideration was placed on retention of the overall protein structure via RMSD of C_α_ atoms. Harmonic restraints on all protein heavy atoms (100 kcal/mol) and fixed constraints on ligand heavy atoms was maintained throughout initial equilibration, which included: heating to 300K with gradual scaling of temperature by 0.2 K every 100 femtoseconds (fs); application of the NPT ensemble (isobaric-isothermal) with Nosé-Hoover temperature control at 300K for several ns until the reported pressure was consistently near 1 atm. Dynamics were propagated with the Leapfrog integrator with a time-step of 2 fs. Van der Waals forces were truncated with a switching function between 11 and 12 Å. Particle Mesh Ewald was used for electrostatic interactions with a real-space cutoff of 12 Å, a κ of 0.333 Å, and a grid ∼1 Å. Protein heavy atom restraints were then gradually released in a radial fashion (side-chains first, then backbone) from a defined point in the catalytic center using the MD program NAMD (v. 2.8) for 10 ns. Final velocities and equilibrated coordinates were used to generate the necessary Anton input files (ark files) using the DESRES provided guesser scripts.

Dynamics in Anton were propagated using the RESPA integrator with a time-step of 2 fs. NPT ensemble at 300K with Berendsen thermostat/barostat was used. Production was done on the 512 node Anton machine with structures collected every 240 picoseconds. All simulation analyses were carried out using Bio3D^25^ in R package. C_α_ root-mean-square deviation (C_α_RMSD), C_α_ root-mean-square fluctuation (C_α_RMSF), and C_α_ radius of gyration (C_α_RGYR) were evaluated to determine the conformational changes over the course of microsecond simulation. Dynamic cross correlation (DCC) and principal component (PCA) were performed to determine the intramolecular “cross-talks” between domains and principal motion observed throughout the simulation.

### Site-Directed Mutagenesis of HepI from E. coli K12

All materials, solvents, competent cells were obtained as previously reported29. The Q5 High-Fidelity PCR kit from New England Biolabs was used according to its instructions to perform mutagenesis using *E. coli K-12* strain MB1760 960 bp HepI gene sub-cloned into pTOM-15b. New England Biolabs thermocycling conditions for routine PCR was used and the amplified DNA was then transformed into XL10-Gold competent cells, which were then incubated in a 5 mL LB/AMP overnight growth for plasmid DNA extraction by miniprep. Once the mutation was confirmed by sequence alignment with HepI wild type DNA and the respective primers for each mutant, the purified plasmid was transformed into BL21-AI cells as per Agilent’s instructions.

P216G Forward Primer: gcgcgccccacccaagtttaatccgtattcctg

P216G Reverse Primer: caggaatacggattaaacttgggtggggcgcgc

P240G Forward Primer: gttgaagtattgggcaagatgagtctggaaggcgttg

P240G Reverse Primer: gttgaagtattgggcaagatgagtctggaaggcgttg

G280P Forward Primer: ggatagacccaatatcacggtttatccgccaaccgatccg

G280P Reverse Primer: cggatcggttggcggataaaccgtgatattgggtctatcc

G288P Forward Primer: ccgggattaattcctgggtatgggaagaatcagatggtatgtagggctcc

G288P Reverse Primer: ggagccctacataccatctgattcttcccatacccaggaattaatcccgg

### Mutant protein expression and purification

All buffers for SDS-PAGE gels and protein purification protocols are the same for as wild-type HepI, as reported in previous literature30. All mutants transformed into BL21-AI including WT were grown and expressed with the same conditions as described previously29. Briefly, cells were harvested, resuspended in 20 mL bind buffer (20 mM HEPES, 1 µM imidazole, 500 mM NaCl, pH=7.5) per liter of growth on ice and lysozyme was added. The cells were then homogenized for 4-8 cycles and the lysate clarified by centrifugation at 13,000 rpm for 1 hour. The lysate was purified via FPLC with parameters described previously. Monitoring of 280 nm was used to identify fractions containing HepI and those fractions were then pooled and concentrated. The concentrated protein was desalted into a storage buffer of 100 mM HEPES, 1 M KCl, pH 7.5 using a BioRad P6 polyacrylamide SEC column. The fractions containing protein were then concentrated and precipitated in an equal volume of saturated ammonium sulfate and stored in amber vials at 4°C.

### HepI Mutant Kinetic Characterization

HepI substrates were prepared as previously described^26^ for kinetic analysis of HepI and its mutants. Steady-state kinetic data was collected using a UV-Vis spectrophotometric technique of a pyruvate kinase-lactate dehydrogenase coupled assay which monitors the reduction of NADH overtime upon release of ADP. Data collected was analyzed using the Michaelis-Menton approximation and reported in Table 1.

**Table 1.** Kinetic constants for the mutant HepI proteins.

| Hepl | $k_{\text{cat}}$ ($\text{s}^{-1}$) | Fold Change | $K_M$ ($\mu\text{M}$) | Fold Change | $k_{\text{cat}}/K_M$ |
| --- | --- | --- | --- | --- | --- |
| Wild type | $0.59 \pm 0.09$ | - | $8.5 \pm 3.6$ | - | $6.8 \times 10^4$ |
| P216G | $0.21 \pm 0.02$ | 2.8 ↓ | $1.6 \pm 0.6$ | 5.6 ↓ | $1.3 \times 10^5$ |
| P240G | $0.19 \pm 0.01$ | 3.1 ↓ | $0.9 \pm 0.3$ | 10 ↓ | $2.0 \times 10^5$ |
| G280P | $0.21 \pm 0.02$ | 2.8 ↓ | $1.5 \pm 0.5$ | 6.0 ↓ | $1.3 \times 10^5$ |
| G288P | $0.48 \pm 0.07$ | 1.2 ↓ | $10.2 \pm 0.4$ | 1.1 ↑ | $4.7 \times 10^4$ |

## Results

The tertiary structural stability of both GT-B systems for the full length of the simulation was assessed by calculating the enzymes’ global C_α_ Root Mean Square Deviation (C_α_RMSD) and Radius of Gyration (C_α_RGYR). The observed C_α_RMSD for both HepI and GtfA are shown in Figure 1. The C_α_RMSD for the two-domain HepIC_α_ HepI is highly oscillating between 1.5 and 3.5 Å, with a dominant decreasing trend for approximately the first 0.6 µs, followed by a sharp, consistent increase for the remainder of the simulation (**Fig. S1**). Of the structural ensemble represented by the full HepI trajectory, 11.5% of the frames report enzyme structures with C_α_RMSD values > 3.0 Å, while 49.4% are between 2.0 Å and 3.0 Å. The C_α_RMSD for GtfA shows a gradual increase in the C_α_RMSD observed over time. A locally weighted scatterplot smoothing (lowess) curve that is fit to the C_α_RMSD data shows the increase to be biphasic: an initial ∼0.75 µs long phase of rapid, steady rise in the C_α_RMSD followed by a final phase represented by a slowly increasing C_α_RMSD cluster centered between 2.5 Å and 3.0 Å. Exactly 1.35% of the structures in the GtfA trajectory report C_α_RMSD values > 3.0 Å, while 78.15% have values spanning from 2.0 Å to 3.0 Å. The C_α_RGYR data of HepI and GtfA continually increase and encompass roughly the same range (∼ 21.0 Å to ∼ 22.7 Å) (**Fig. 2 A&D**). In both enzymes, the C_α_RGYR spread narrows with passing time, gradually concentrating towards higher values. Coupling of the respective C_α_RGYR and C_α_RMSD data for each enzyme highlights a positive correlation between the two variables, although in GtfA, said correlation is evident only after ∼0.4 – 0.5µs. The data suggest that changes in the compactness of both enzymes are connected to the underlying global structural modifications as measured by the C_α_RMSD.

**Figure 2.**
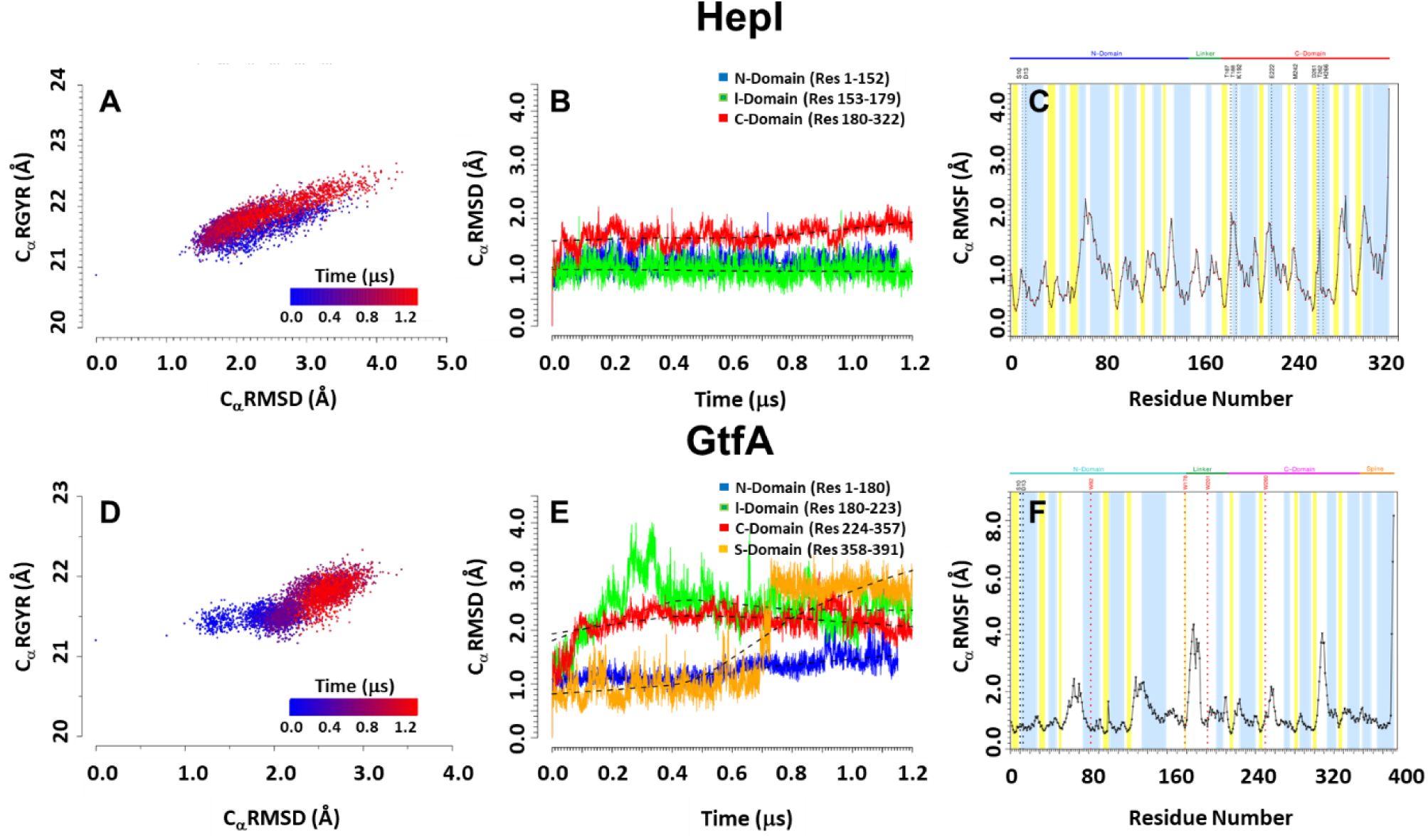
C_α_RGYR, C_α_RMSD and C_α_RMSF plots of HepI (A-C) and GtfA (D-F). **A & D)** The C_α_ radius of gyration (C_α_ RGYR) against for C_α_RMSD are shown to highlight the changing of conformation population over the course of MD simulation from 0.0 (blue) through 1.2 µs (red). **B & E)** C_α_RMSD by each of the glycosyltransferase domain. Black dotted lines represent lowess curves for each domain. **C & F)** C_α_RMSF plot with secondary structures, α-helix and β-sheet regions, highlighted in blue and yellow, respectively.

To explore whether the data are representative of structural instability and/or early microsecond dynamic motions (e.g., loop restructuring, domain repositioning), the structural stability and/or dynamic rearrangements of the ***N*, l**inker, ***C***, and **s**pine (GtfA only) domains were examined (**Fig. 2 B&E**) by calculating: 1) the C_α_RMSD of each domain individually (per-domain: *N*-, *l*-, *C*-, and *s*-C_α_RMSD), and 2) the C_α_RMSD of the grouping by type of each of the domains respective secondary structural elements ([Domain]_α/β_-ssC_α_RMSD). The data for each of these localized C_α_RMSD analyses were calculated after a full MD trajectory to reference crystal structure coordinate alignment was performed that only included the protein regions of interest. The resulting data report exclusively on the changes to the structure of the chosen protein segment(s). As such, the per-domain C_α_RMSD analysis functions as a macroscopic evaluation of the domain’s total structure, while the per-secondary structure type C_α_RMSD serves as a microscopic assessment that focuses on changes to the underlying architecture of the domains.

The average HepI *C*_α_RMSD and ssRMSD for each domain fall under 2.0 Å and display a near-linear trend, with both the *N* and *C* domain α-sheet cores consistently having lower value, less fluctuating ssRMSD data than the enveloping alpha helices. Unlike the HepI *N* and linker domains, the HepI *C* domain is in a higher RMSD value range and steadily increases after ∼0.6–0.7 µs (**Fig. 2B**). The pattern observed in the *C*_α_RMSD beyond ∼0.6 µs is both homologous to that of the C_β_-ssRMSD and matches the rise in the C_α_-ssRMSD during the same time (**Fig. S2**). HepI ssRMSD *N* and *C* domain β-sheet cores consistently having lower average and less fluctuating ssRMSD data than the enveloping α helices. The *N*- and *l*-RMSD are congruent and fit linearly to a lowess curve. On the other hand, with the exception of the *N* domain, increase variance is observed in the per-domain C_α_RMSD GtfA. As in HepI, the C_α_ssRMSD data all fall below 2.0 Å. The GtfA *N*-C_α_RMSD has a more constant, near-linear trend that begins to increase at ∼0.6µs. This sudden increase concurs with the start of the inward displacement of the C-terminal end of N-βIII (res: 50-52) towards the more central N-βII (res: 30-35), as reported in the clearly defined jump at ∼0.6 µs in the N_β_-ssRMSD. The start of a slight increase in the N_α_ssRMSD is also observed at ∼0.4 µs. This corresponds to the outward motion of residues 134-149, which comprise the N-terminal segment of N-αV (res: 134-159).

GtfA *s*-C_α_RMSD initially shows a moderately fluctuating section that quickly shifts at ∼0.7 µs to a tighter, slowly decreasing segment in a higher C_α_RMSD value range (**Fig. 2E**). Frames at the transition time point clearly portray an unwinding of the protein’s last three C-terminal residues (res: 389-391 on s-αII), with no alterations or significant displacement to the rest of the spine’s backbone structure. GtfA *C*- and *l*-C_α_RMSD result in a near equivalent logarithmic dataset peaking at ∼0.2 µs and ∼0.3 µs, respectively, followed by a period of steady decline. The C_α_-, C_β_-, or l_α_-ssC_α_RMSD data do not detail any significant structural changes to account for the shape of the *C*- and *l*-C_α_RMSD data (**Fig. S1**). With the exception of the global C_α_RMSD, none of the domain or ss-RMSD analyses consider loop (lp) or 3_10_-helix regions. From the C_α_ Root Mean Square Fluctuation (C_α_RMSF), the largest time-averaged atomic fluctuations (C_α_RMSF > 2.0 Å) in both enzymes primarily correspond to loop, 3_10_, and adjacent N/C-terminal secondary structure residues (**Fig. 2 C&F**). In HepI, these areas are: 63-70 (N-α3 C-term [63-64], N-lp5 [65-67], N-αIV N-term [68-70]), 136-137 (N-lpX), 188-191 (C-lpI C-cap [188], C-3_10_I [189-191]), 218-221 (C-lpII C-cap [218], C-αII [219-221]), 263 (C-αIV N-cap), 280-285 (C-lpVII), 300-303 (C-3_10_III), and 321-322 (C-αV C-term). In GtfA, the residues are: 64-65 and 70 (N-lpII), 127-128 (N-3_10_IV), 183-193 (l-lpI), 266 and 268 (C-lpIII), 315-322 (C-lpVII), and 389-391 (C-αVII C-term). The locations of these high C_α_RMSF regions on the enzymes’ structures generally correspond to the side boundaries of the binding cavity on the *N* and *C* domains. The equivalence is solely spatial and does not always correlate to similar structural elements. A larger proportion of the high RMSF areas are positioned on the catalytic side of the cavity, primarily encompassing regions in the vicinity of the volume occupied (HepI) or predicted to be occupied (GtfA) by the sugar moiety of the co-factor. β-sheets in both systems consistently have the lowest C_α_RMSF values (< 1.0 Å) and are data trough points. The α-helices demonstrate more fluctuation, with the high C_α_RMSF points corresponding to the helical N/C-terminal end residues.

The dynamic motions of both systems are not limited to just loop and 3_10_-helix movements, as reported in the principal component analyses (PCA) (**Fig. 3&S3**). In the HepI and GtfA simulations, 60.1% and 50.5% of the overall structural variance lies in the first three principal components, respectively. To better describe the underlying structural motions contained in each PC, only the extreme points that sit on the principal component (PC) axes were considered. Each point is a single trajectory frame, whose protein backbone arrangement (C_α_ coordinates) represents the endpoint (end structure) of the pure structural motion comprised in each PC. Overlay of both extreme points for each PC generates a structural grouping, which accounts for the full dynamic range within each PC.

**Figure 3.**
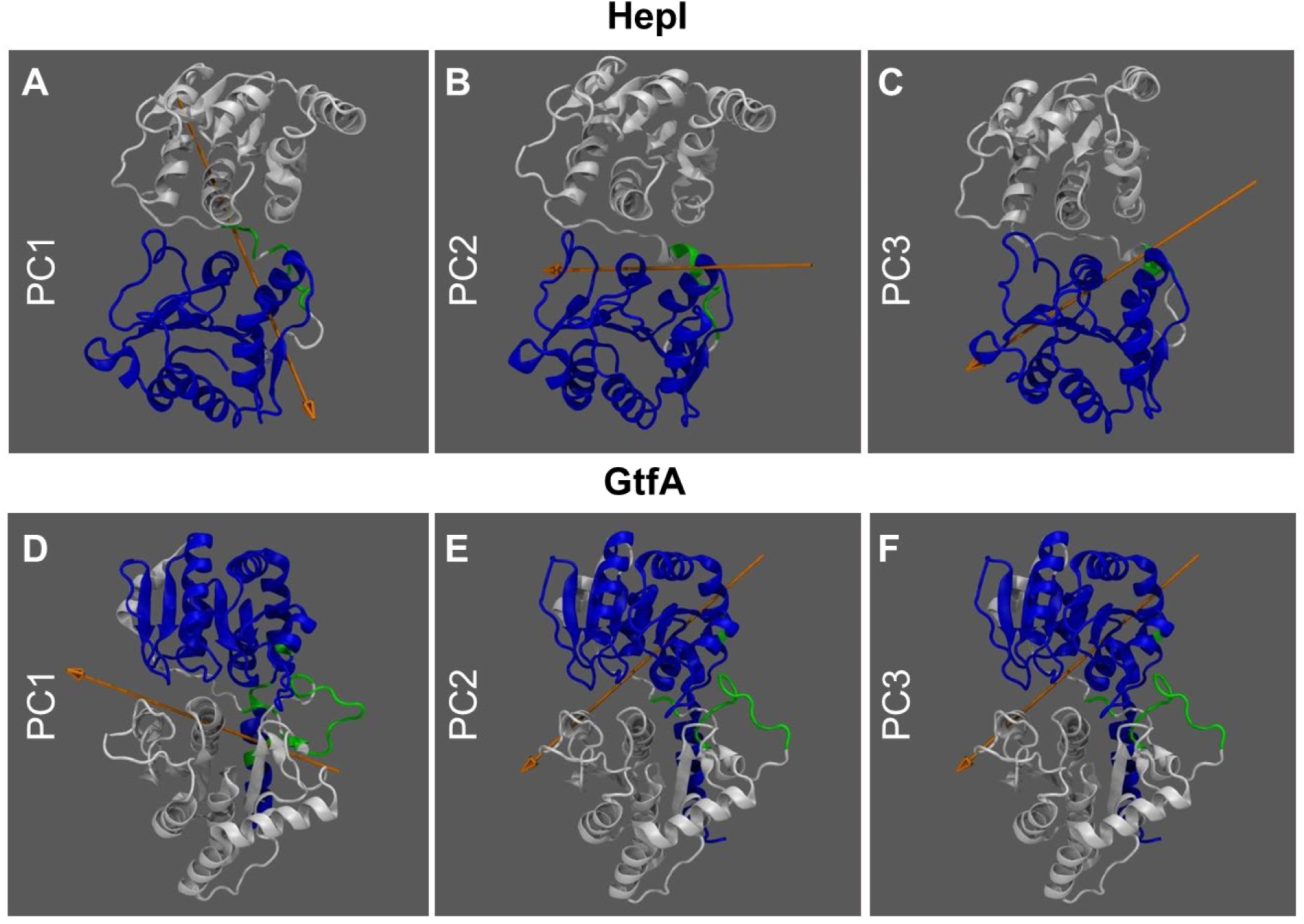
Top 3 principal dynamic motions of HepI (A-C) and GtfA (D-F) derived from MD simulation. HepI and GtfA domains, C-domain, N-domain, and linker region are represented by white, blue, and green ribbons, respectively. The top three components, accounting for 60.1% and 50.5% of dynamic motions, are plotted separately for both HepI (top) and GtfA (bottom). Orange arrow represents direction of rotational axis.

PC1 of HepI (PC1_H_, 37.5% structural variance), pertains to the ∼24.2°-33.1° lateral, outward cork-screw rotation of the *C* domain about the *N* domain, hinged around linker residues 164-166 and 168-174 of l-lpII (**Fig. 3 A-C**). The rotational axis diagonally intersects both domains, from the lateral edge of catalytic side of the *N* domain to the lateral edge of the non-catalytic portion of the *C* domain. PC2_H_ (12.7% structural variance), accounts for the ∼21.6°-27.9° clamp-like motion of the *N* and *C* domain, hinged around residues 168-175 of l-lpII (**Fig. 3B**). The PC2_H_ axis of rotation lies parallel to the longest plane of the inter-domain cavity, expanding from the peripheral edges of the catalytic and non-catalytic areas of the binding cavity. PC3_H_ (9.95% structural variance), involves a ∼13.9°-14.7° rocking motion of the *N* and *C* domain, whose rotation axis diagonally crosses the mid-point of both domains, extending from the back-end to frontal face of the binding cavity (**Fig. 3C**). The motion hinges about linker (164-165 [l-lpI], 166-169 [l-αI]) and *C* domain (180-184 [C-β1], 262 [C-βIV], 263 and 266-267 [C-αVI], 282-284 and 288-291 [C-lpVII]) residues.

Of all the HepI PC transitions, the most prevalent occur in PC1_H_ (cork-screw rotation) and PC3_H_ (rocking motion). Sequential examination of the PC_H_ data by time-clusters reveals the enzyme system progressing initially from octant II (O-II) to O-I (-PC1 → +PC1) (**Fig. 4 A&B**). The starting structural configuration of HepI is in O-II (-PC1, +PC2, +PC3). Octant II accounts for 13% of the total trajectory ensemble and is not only occupied predominantly by structures from the first 80 ns, but also sporadically by later structures in the simulation. Trajectory frames that lie in O-II are presumed to have structural configurations like that of the reference crystal structure. The succeeding transitions are: O-I to O-V (+PC3_H_ → -PC3_H_) and O-V to O-VI (+PC1_H_ → -PC1_H_). It is after the latter transition that the system has a defined change in PC2_H_ (clamp motion) by going from O-VI to O-VII (+PC2_H_ → -PC2_H_). The subsequent shifts predominantly take place in –PC2_H_ space and are: O-VII to O-III (-PC3_H_ to +PC3_H_), O-III to O-IV (-PC1_H_ to +PC1_H_), and O-IV to O-VIII (+PC3_H_ to –PC3_H_). There are, however, single points or segments of the trajectory that deviate from the above described progression and are found in an octant other than that of the time-cluster it would be affiliated with.

**Figure 4.**
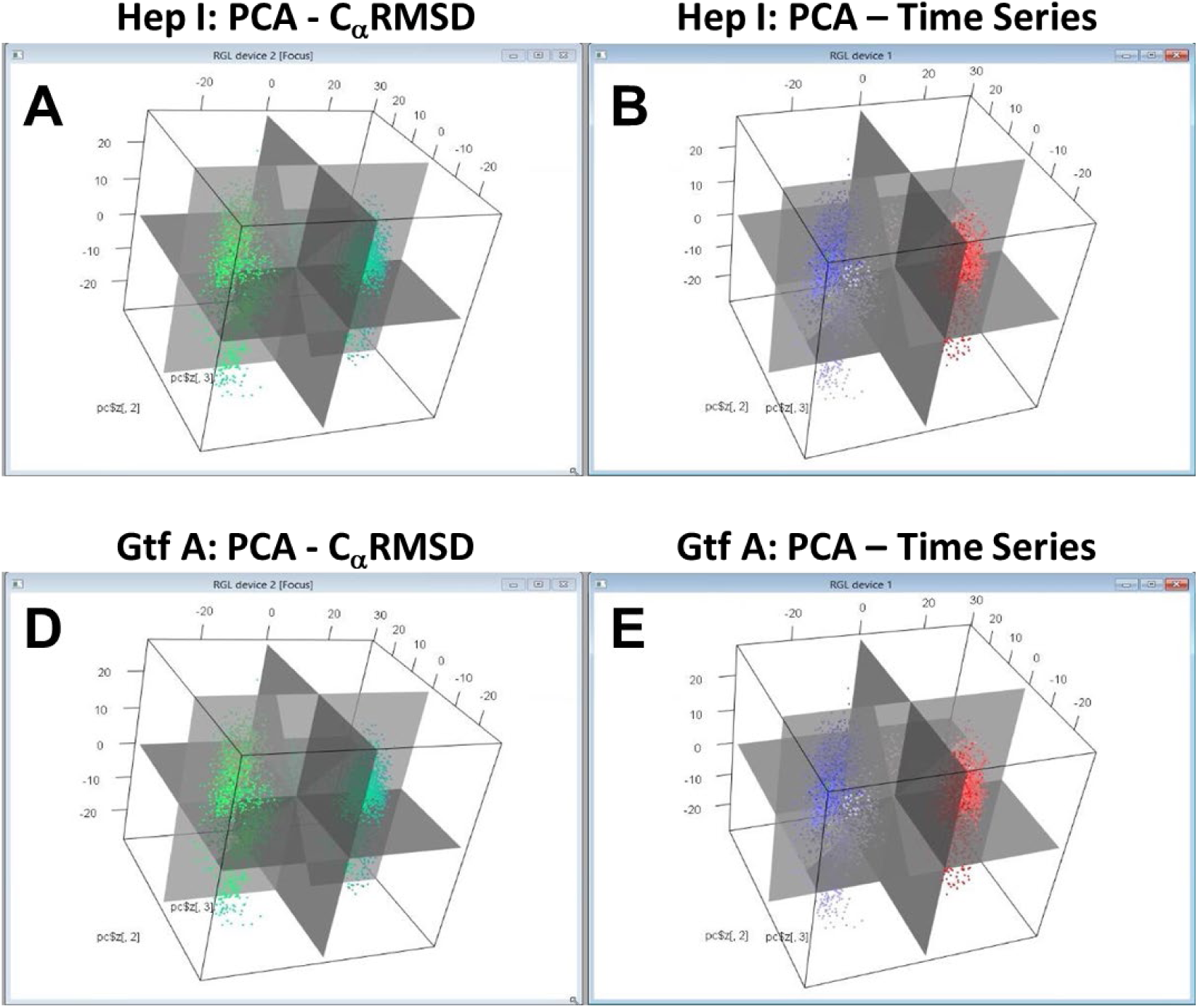
Conformational distribution of the top three principal components (3D-PCA) plotted against RMSD (green to teal) and time from 0.0 (blue) through 1.2 µs (red).

In PC1 of GtfA (PC1_G_, 33.5% structural variance), the motion is akin to the frontal and posterior clamp-like movement of the *N* and *C* domain observed in PC2_H_ (**Fig. 3 D-F**). The PC1_G_ rotational axis also lies in the inter-domain cavity and spans the binding pocket from edge to edge. The degree of rotation about the axis is ∼12.7°-15.6°, hinged on specific linker (177-179 [N-β6], 180-201 [l-lpI]) and spine residues (369-370 [s-3_10_I], 373-374 and 379-380 [s-αII]). PC2 of GtfA shows a ∼11.0°-13.8° rotational displacement of the *C* domain about the *N* domain, hinged on select linker (181-194, 196-197, and 201-202 of l-lpI) and spine residues (372-376 and 378-379 [s-αII]) (**Fig. 3E**). The movement in PC2_G_ is similar to the motion represented by PC1_H_ (cork-screw rotation). PC3_G_ displays the same rocking type motion as in PC3_H_. The PC3_G_ axis of motion also diagonally crosses both domains from the back (catalytic side of the *N* domain to the front (non-catalytic side of the *C* domain) faces of the binding cavity (**Fig. 3E**). The motion is a ∼11.0°-13.8° rotation, hinged on linker (184-194 and 196-197 [l-lpI]) and spine (374-379 [s-αII]) residues.

The prevailing PC_G_ transitions occur between PC2_G_ (cork-screw rotation) and PC3_G_ (rocking motion). As in HepI, the starting GtfA structure also lies in O-II, which includes all frames from the first 0.3µs of simulation (18% of the total trajectory ensemble). By tracking the evolution of PC_G_ data clusters over time, the system’s progression is: O-II to O-III (+PC2_G_ → -PC2_G_), O-III to O-VII (+PC3_G_ → -PC3_G_), and O-VII to O-VI (-PC2_G_ → +PC2_G_) (**Fig. 4 C&D**). After the second cork-screw rotation event, there is a clear transition from O-VI to O-V (-PC1_G_ to +PC1_G_). Note that the sequence of transitions that lead to the shift in the clamp-like PC is the same in GtfA (PC1_G_) and HepI (PC2_H_). Unlike HepI, there are no obvious structures that are found in –PC1_G_ space after the transition takes place. Additionally, the subsequent steps (type of motions) correlate in order with those seen in HepI: O-V to O-I (-PC3_G_ to +PC3_G_), O-I to O-IV (+PC1_G_ to –PC1_G_), O-IV to O-VII (+PC3_G_ to –PC3_G_). Another commonality between the two enzymes is the presence of structures in octants other than that expected, given its corresponding time value.

Examination of the dynamic cross correlation matrices (**Fig. 3**) shows that residues near each other in their primary sequence demonstrate strong positively correlated motions, as illustrated by the deep red coloring on and near the diagonal of the matrix for both HepI and GtfA. Additionally, off-diagonal regions within the same domain predominantly demonstrate strong positively correlated motions, while off-diagonal regions that represent the relationship between the two domains (i.e. HepI N-terminal domain residues 1-152 with C-terminal domain residues 180-322) demonstrate predominantly negatively correlated motions. Specific examination of correlations to residues in HepI that are known to participate in conformational changes associated with binding of the sugar acceptor ligand (including Lys 64, Arg 63 and Arg 120),^15, 26-27^ identified a series of glycine and proline residues distant from the binding sites of either substrate (>25 Å away from the binding site of the sugar acceptor but within 15 Å of the sugar donor binding site) with strong negative correlated motions (Pro 216, Pro 240, Gly 280 and Gly 288). Each of these residues was individually mutated, with prolines being introduced where the protein originally had a glycine, and vice versa to yield mutant proteins with the greatest possible perturbation of their conformational flexibility at each position. The resultant proteins were then expressed, purified and kinetically characterized using high concentration of the sugar donor substrate (ADP-L-glycero-D-manno-heptose) and varying concentrations of the sugar acceptor substrate (ODLA) to assess the impact of these residues on its binding (Table 1). Each of the mutant HepI proteins exhibited mildly diminished *k*_cat_ values (with reductions of 3.1-fold as compared to wild type HepI, or less). Notably, three of the mutants P216G, P240G and G280P all had reductions in *K*_m_, resulting in Michaelis constants of 1.6, 0.9 and 1.6 µM, respectively, which lead to an overall enhancement in catalytic efficiency relative to wild type HepI.

## Discussion

We demonstrate that simulating the GT holo to apo transitions led to trajectories that encompass a robust sampling of conformational substates in both enzymes. We perform MD statistical analyses and employ 3-D principal component analysis (PCA) to highlight structural retention (global and by domain) during the course of the simulation, as well as identification of the conformational substates and their transitions along the holo to apo pathway. We show these substates to correspond to domain flexibility (domain repositioning) events that depict the same hierarchy in both enzymes regardless of MD starting state (GtfA: closed holo-ternary; HepI: open holo-binary). Our observation of a hierarchy of substates is akin to similar processes reported via *in-vitro* and/or *in-silico* studies in myoglobin ^28-29^ and in adenylate kinase.^30^ In those systems, the enzymes transition between substates with preferred directionality rather than by stochastic sampling. Whether the reported conformational directionality extends to familial enzymes of those systems remains to be fully determined.

For HepI, the predominance of per-domain RMSD values ≤ 2.0 Å highlight the structural permanence and stability of the domains. Structurally, the changes to the *C* domain starting at ∼0.6 µs correspond to a slight outward displacement of the helical N-termini (N-cap and adjacent neighboring residues in both directions) of α-helices C-αII (res: 219-231), C-αIII (res: 243-253), and C-αIV (res: 262-271). The N-termini of all three helices form part of the binding cavity. Specifically, the latter helix (C-αIV) is positioned at the cavity mid-point, directly over the enzyme’s catalytic core on N-βI, while the N-termini of the two former helices are situated on the non-catalytic side of the binding cavity. In the crystal structure (PDB: 2H1H), these three regions associate with the adenine base of the sugar co-factor. Residue E222 of C-αII forms hydrogen bonds with the 2’ and 3’ hydroxyls of the ribose. Residue M242, which is the amino acid right before the start of the N-terminal end of C-αIII, forms a hydrogen bond between its backbone carbonyl oxygen and the amine of the adenosine base. In C-αIV, the N-cap T262 hydrogen bonds to an α-phosphate oxygen of the diphosphate group. The same small outward motion is also evident from ∼0.6 µs onward in C-βV (res: 275-279) and C-βVI (res: 294-298). Both β-sheets are located on the catalytic side of the binding cavity.

Ultimately, despite obtaining 1.2 µs MD simulations of both enzymes, we were only able to observe the partial interconversion between the open and closed conformations that provided a glimpses into the principle motions along the path for catalysis (**Fig. 3&4**). Despite not realizing our full goal, the resulting trajectory data enabled us to (i) examine the root mean square fluctuation (RMSF) of residues, (ii) assess the dynamic cross correlation (DCC) of the protein movements and (iii) identify the principal components of the motions of both HepI and GtfA. We noticed dramatic evolution of the C_α_RMSD for GtfA (closed with substrates removed) and modest changes in HepI (open apo), consistent with our hypothesis that the presence of substrate induces changes in the protein conformation which would be reversed after removal of the substrates. The C_α_RMSF analyses enabled the identification of dynamic loop regions of both proteins that were adjacent to the ligand binding sites that were suspected to be essential for ligand induced conformational changes. Subsequently, both the 60s and 120s loops of the N-terminal domain (residues 1-152) of HepI were shown to be critical for substrate binding experimentally, which has also been subsequently confirmed by crystallography.^2-3, 31^

The DCC analyses (**Fig. 5**) helped reveal that both proteins have a high degree of negatively correlated motions between the two domains, consistent with the enzyme practicing the open-to-closed motions even in the absence of sufficient time to perform the complete interconversion of those two states. Examination of the DCC matrices, specifically at the 60s and 120s loops identified above, revealed that there were numerous proline and glycine residues within the HepI C-terminal domain (residues 180-322) that exhibited negatively correlated movements with these ligand binding residues. Since glycine and proline are known to exhibit dynamics that are distinct from other amino acids due to the lack of substitution at C-α (as with glycine) or the cyclical nature of the substitution at Cα (as with proline), and because others have noted their involvement in protein flexibility and protein folding^32-33^, we mutated these residues to test the importance of their conformational variability on the overall behavior of the protein. We hypothesized that the rearrangement of the 60s and 120s loops might communicate substrate occupancy of the N-terminal domain to the C-terminus, and recent mutagenesis of these residues (where Glycines were mutated into Prolines and vice versa; (**Table 1)** yielded proteins that bind 10x more tightly to the sugar acceptor to the N-terminal domain. The most pronounced effect was observed in the P240G mutant, which had nanomolar binding affinity for the sugar acceptor substrate, ODLA, and an overall catalytic efficiency of 2.0 × 10^5^. P240G and the other mutated residues are all >25 Å away from the ligand binding site for the sugar acceptor, yet their mutation to alter their conformational lability enhances the enzyme’s ability to adopt the Michaelis complex by an as yet unknown mechanism that requires further exploration. This finding in HepI supports the conclusions by Warshel and coworkers^34-35^ that protein dynamics are primarily responsible for assisting the enzyme in adopting a state where chemistry can occur, and not in yielding a dynamical rate enhancement. Lastly, the analysis of principle components 1-3 of both HepI and GtfA (**Fig. 3&4**) revealed that despite starting in different conformations (HepI open and GtfA closed), the two proteins maintain the same primary principle components and have a shared conformational hierarchy. This suggests to us that the overall dynamics of these proteins are a conserved feature of the GT-B structural scaffold, even when the proteins evolutionarily originate from different ancestry with little to no sequence similarity and catalyze reactions with the opposite stereochemical outcomes.

**Figure 5.**
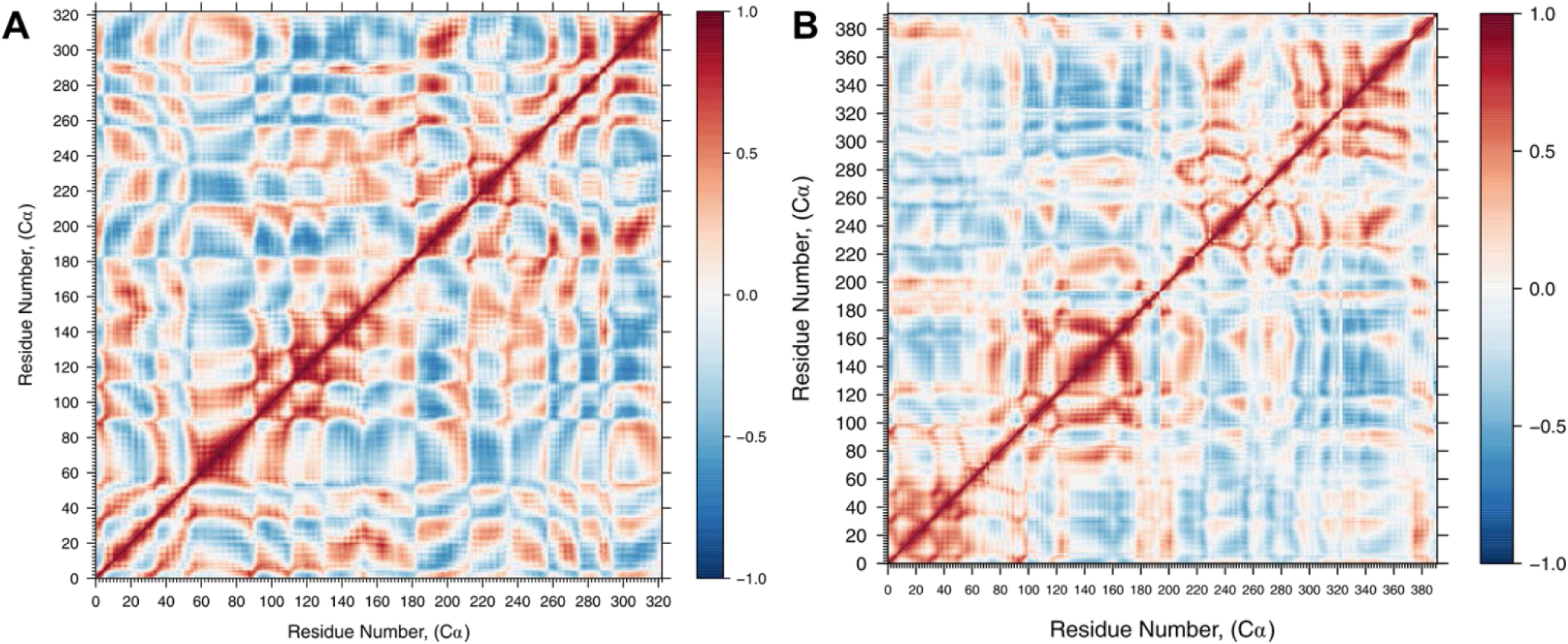
Dynamic C_α_ residue cross-correlation (DCC) of residue fluctuation over 1.2 µs MD simulation showing positively correlated (red), negatively correlated (blue), and uncorrelated (white) regions of **A)** HepI and **B)** GtfA. Black boxes denote 60s and 120s loop regions identified for mutagenesis.

## Conclusion

This is the first ever microsecond MD simulation of GT-B glycosytransferases, and the study suggests that C_α_RMSD by itself is not a good test to determine structural stability at higher timeframes, but rather the protein simulations need to be examined for each individual domain and require more in depth C_α_RMSD analysis that looks at the individual structural constituents and their structural evolution over time. The simulations in these two evolutionarily unrelated, but structurally conserved glycosyltransferases has given us insight into how the structural scaffold itself, and not the primary amino acid sequences are controlling the protein dynamic modes. HepI and GtfA which catalyze reactions with opposite stereochemical courses (specifically HepI catalyzes a reaction with an overall inversion of stereoconfiguration at the anomeric position while GtfA catalyzes a reaction where the anomeric stereoconfiguration is retained) further suggests that the protein’s structure itself is designed to stabilize the formation of the conserved oxocarbenium ion intermediate. That the conformational directionality and the chemical environment of the active site is a preserved feature of these structures, and is encoded in the fold rather than the sequence perhaps illustrating why convergent evolution of this structural fold has been recapitulated to produce a series of enzyme catalyzing sugar transfer reactions. Additionally, the observation that mutagenesis of these distal non-ionizable residues which exhibit correlated motions to those residues that are important for ligand binding, suggests that the enzyme dynamics for HepI are enabling the preordering of the enzyme into the Michaelis complex and not chemistry. While dynamic motions are often considered to be critical for traversing activation energy barriers, in this case, the mutagenesis data is consistent with the theory that dynamic modes are enabling the electrostatic organization of the Michaelis complex and not making a significant contribution to catalysis.

## Abbreviations

GT: glycosyltransferase
HepI: heptosyltransferase I
heptose: L-glycero-D-manno-heptose
ADP-heptose or ADPH: ADP-L-glycero-D-manno-heptose
LPS: lipopolysaccharide;
ODLA: O-deacylated *E. coli* Kdo2-lipid A;
Trp: tryptophan;
Pro: proline;
Gly: glycine;
Arg: arginine;
Lys: lysine;
CD: circular dichroism;
MD: Molecular Dynamics;
PDB: Protein Data Bank
PCA: Principal Component Analysis,
DCC: Dynamic Cross Correlation

---

**Figure S1.**
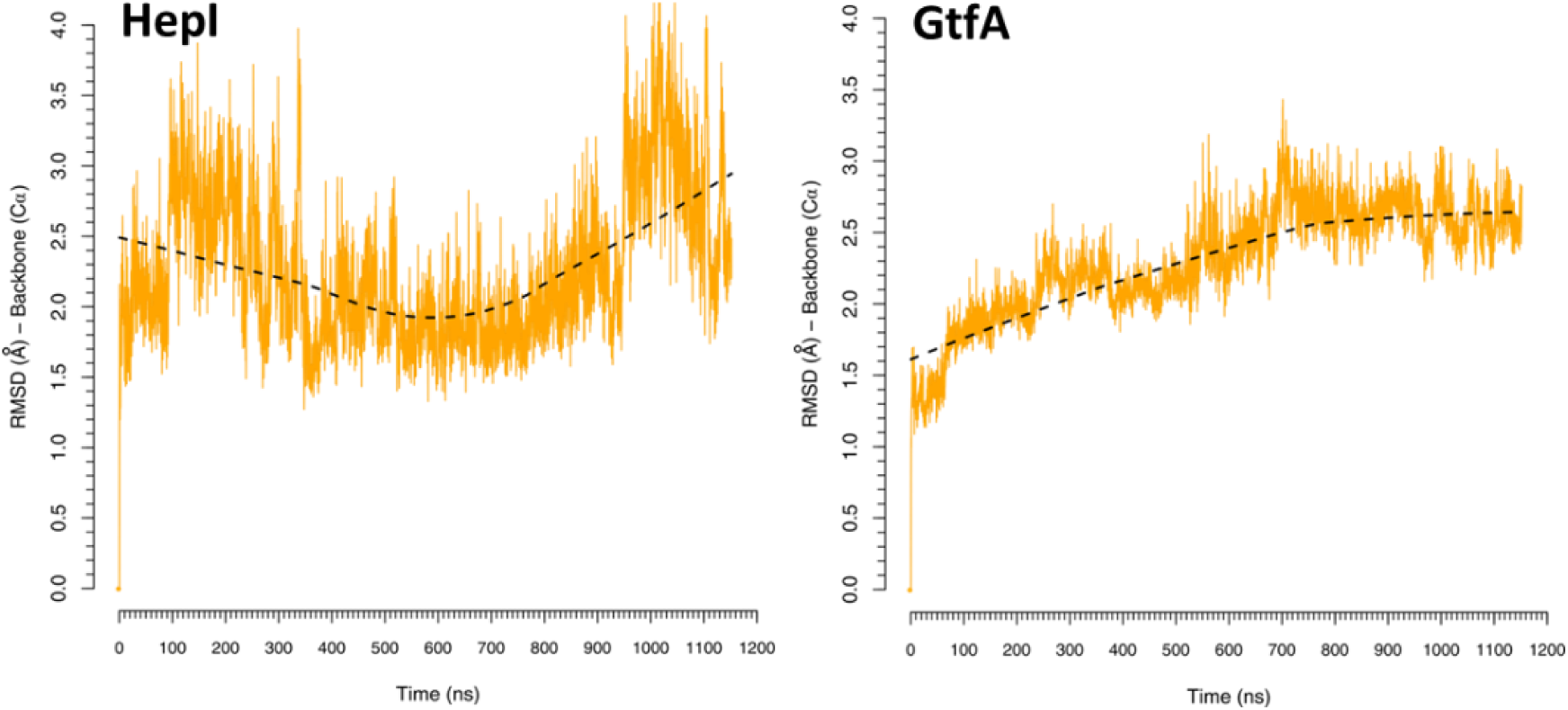
C_α_RMSD plots of HepI and GtfA.

**Figure S2.**
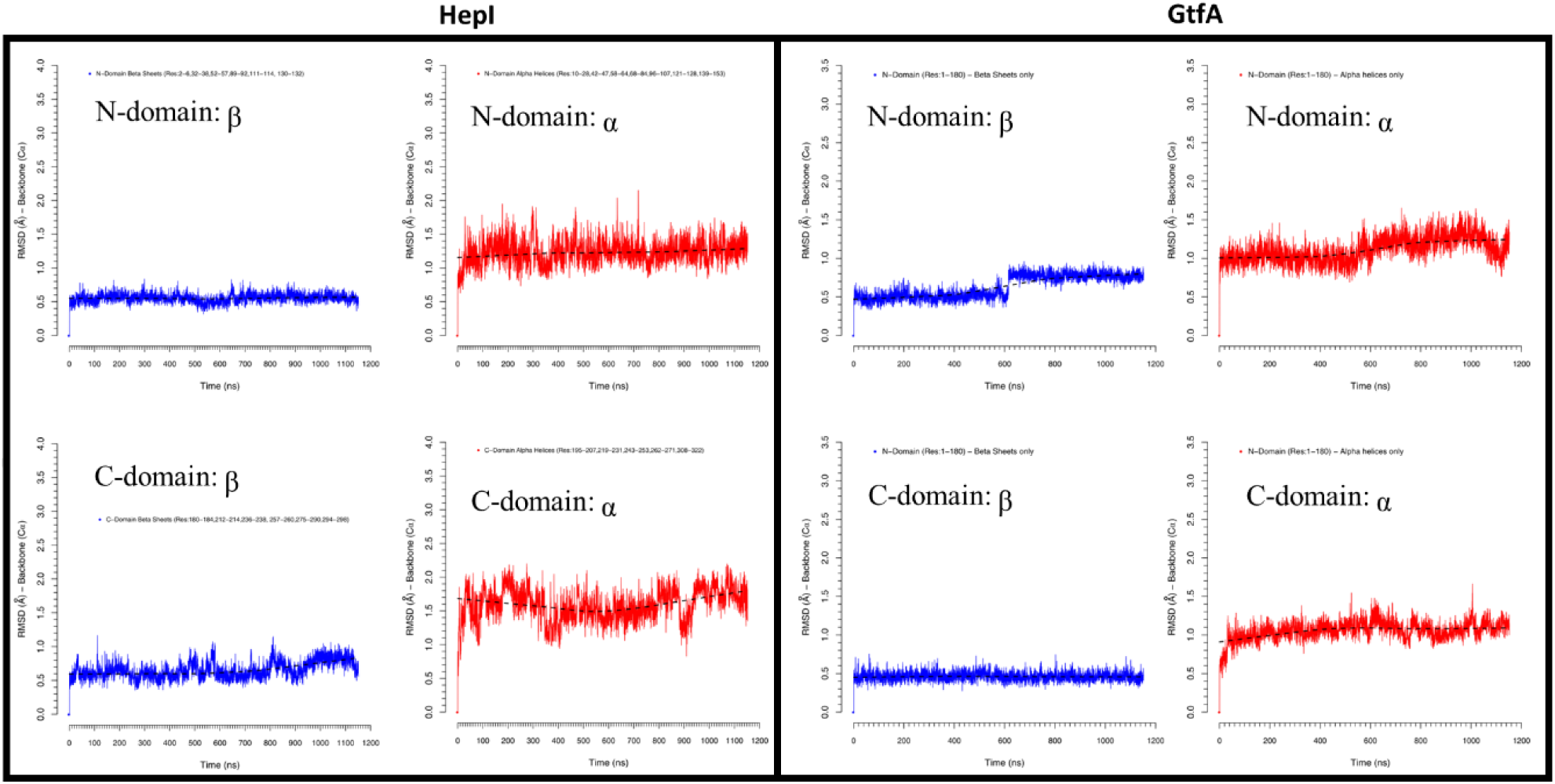
C_α_RMSD of Rossman fold motif secondary structure elements in HepI and GtfA from MD simulation.

**Figure S3.**
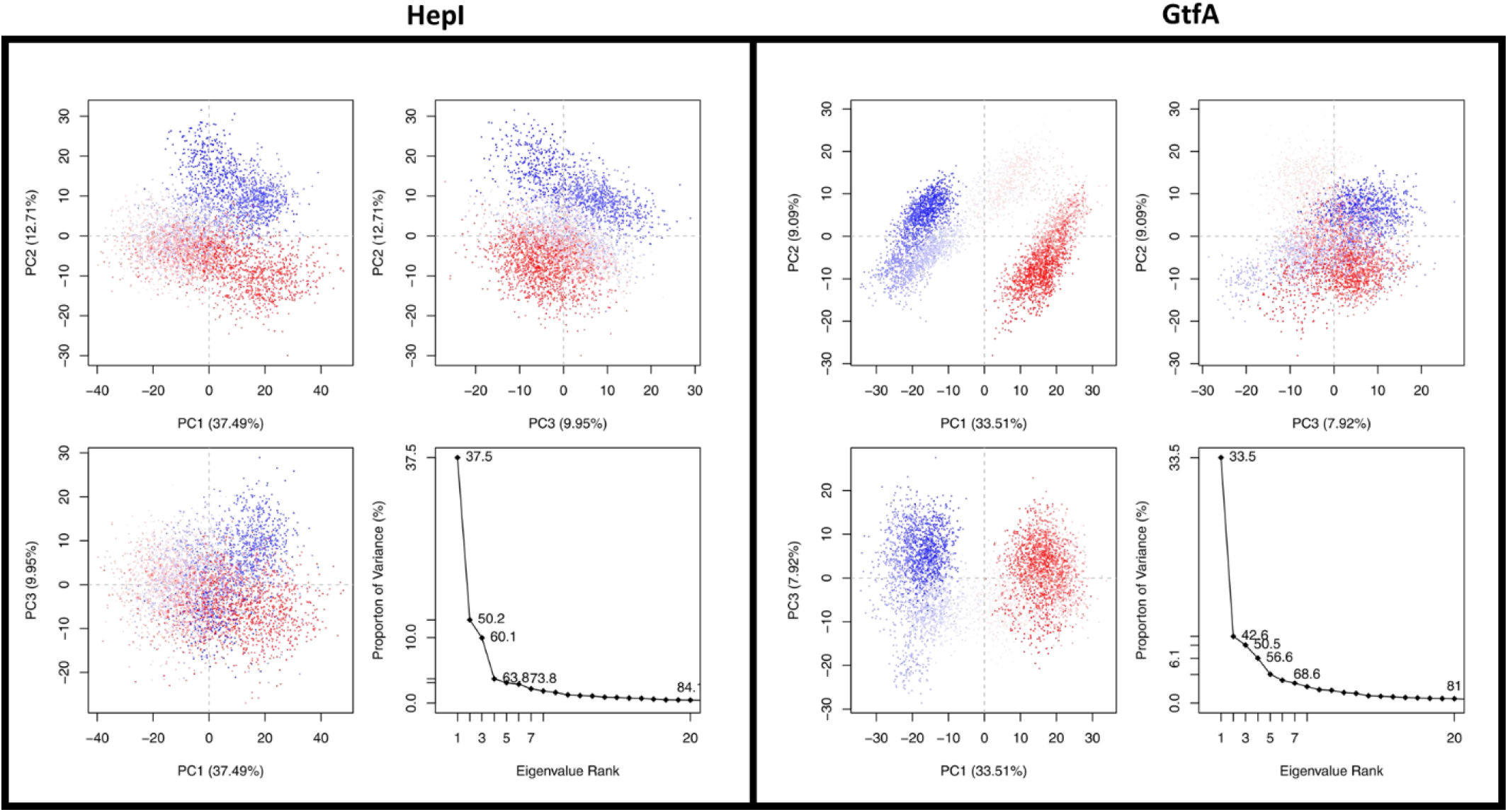
PCA scatter plot of first three PCs against each other and percentage proportion of variance of each PC for HepI and GtfA

